# Transgenerational effects increase the vulnerability of a host-parasitoid system to rising temperatures

**DOI:** 10.1101/2024.08.16.608228

**Authors:** Natalie L. Bright, Jinlin Chen, J. Christopher D. Terry

## Abstract

1. Transgenerational effects, non-evolutionary processes by which environmental conditions in one generation influence the performance in subsequent generations, are hypothesised to have substantial consequences for population dynamics under stochastic environments. However, any direct apparent detriment or advantage these processes generate for a focal species may be counteracted by concurrent effects upon interacting species.
2. Using an experimental *Drosophila*-parasitoid model system, we determined how the previous generation’s thermal environment impacts the thermal performance of both hosts and parasitoids. We found substantial responses in both trophic-levels, with potential evidence for both condition-transfer effects and adaptive transgenerational plasticity.
3. We used these results to parameterise discrete-time simulation models to explore how transgenerational effects of thermal conditions and temporal autocorrelation in temperature are expected to impact the time to extinction for this host-parasitoid system under climate change. The models predicted that transgenerational effects would significantly hasten the time to extinction, largely through a reduction in estimated average performance. Under the assumptions of one of the population dynamics models trialled, we identified an additional hastening of extinction from the combined effect of both host and parasitoid transgenerational effects.
4. Our research demonstrates how community-level consequences of transgenerational effects may impact a population’s sensitivity to climate change under a fluctuating environment and highlights the need to quantify and contextualise thermal transgenerational effects in their ecological setting.

## Introduction

Transgenerational effects, where the conditions experienced in earlier generations impact the performance in subsequent generations without evolutionary change (Galloway & Etterson, 2007), are increasingly recognised as an influential determinate of climate change risk (Donelson et al., 2018). Where there is autocorrelation in environmental fluctuations facing successive generations (Heino et al., 2000; Petchey et al., 1997), adaptive transgenerational effects may help populations to persist, or alternatively could also shorten persistence if negative impacts accumulate through condition-transfer. However, the impact of transgenerational effects in a multispecies context has been underexplored (Malinski et al., 2024) despite it being widely appreciated that direct effects of climate change on species will be modulated by the wider community context (Carroll et al., 2024; Chen & Lewis, 2023; Davis et al., 1998; Gilman et al., 2010; Tylianakis et al., 2008; Urban et al., 2016). The impact of transgenerational effects in a focal species response may be offset or exacerbated by corresponding responses of interacting species.

Transgenerational effects can include both condition-transfer and adaptive plasticity effects through anticipatory effects, which can be challenging to disentangle (Sánchez‐Tójar et al., 2020). Condition-transfer, the direct modulation of the amount of investment the parental generation can make in the next, is likely the most widespread form of transgenerational effect (Bonduriansky & Crean, 2018). Anticipatory effects, on the other hand, suggest that parents are able to actively influence their offspring’s phenotypes to suit the environment anticipated to experience by the next generation (Burgess & Marshall, 2014). Such priming could reduce the lag between environmental signal and phenotypic response (Cavieres et al., 2020) when environmental fluctuations are predictable, thus significantly altering the extinction risk posed by climate change (Salinas & Munch, 2012). However, the supporting evidence for prevalence of transgenerational effects is still relatively weak (Sánchez‐Tójar et al., 2020; Uller et al., 2013) and the magnitude and direction context-dependent (Baker et al., 2019; Yin et al., 2019). Furthermore, identifying a transgenerational effect is only the first step to demonstrating its impact – the effect must be contextualized into an environmental (Burgess & Marshall, 2014) and biotic context (Walther, 2010).

Here we investigate the impact of transgenerational effects in a model experimental *Drosophila* host-parasitoid system. Insects are likely to be amongst the most responsive taxa to climate change (Harvey et al., 2023). Parasitoids are ectotherms whose larvae develop by feeding in or on the bodies of other arthropods, representing ∼10% of described insect species (Eggleton & Belshaw, 1997; Godfray, 1994). Many of the processes intrinsic to parasitoids’ life-history demonstrate temperature dependence; for instance, foraging traits, such as successfully locating and ovipositing in a host egg, and attack traits, such as evading the host’s immune defences (Hance et al., 2007; Synodinos et al., 2021) and various responses to fluctuating temperature have been documented in parasitoid attack rates (Costaz et al., 2023; Ismaeil et al., 2013; Le Lann et al., 2021). The potential impact of transgenerational effects on insect fitness has long been acknowledged (Mousseau & Dingle, 1991) and has been demonstrated in *Drosophila* in a variety of contexts (Cavieres et al., 2020; Huey et al., 1995; Schiffer et al., 2013). By strongly regulating arthropod host population dynamics and coexistence between different host species, parasitoids play important roles in maintaining biodiversity and influencing community structure (Godfray, 1994; Terry et al., 2021). Host-parasitoid interactions thus represent an ecologically important subset of biotic interactions, yet are relatively underrepresented in literature on biotic responses to climate change (Furlong & Zalucki, 2017; Hance et al., 2007; Jeffs & Lewis, 2013; Thierry et al., 2019). Host-parasitoid interactions offer an experimentally amenable window to start to parse ecosystem complexity. Since the reproduction of parasitoids is closely linked to their success finding and attacking hosts (Hassell, 2000), the dynamics between host and parasitoids populations are tightly coupled with the high potential for emergent effects to arise in their response to climate change (Furlong & Zalucki, 2017; Thierry et al., 2021).

The temporal autocorrelation of the environmental variable in question is central to the impact of transgenerational effects on population dynamics. Without autocorrelation, if the environment faced by a particular generation is not at all predictable from the previous generation, transgenerational effects can be well-summarised by a long-term average. One route to isolate the impact of considering transgenerational effects is to identify emergent effects on population dynamics with and without autocorrelation. However, identifying an emergent impact on population dynamics is complicated by the fact that autocorrelation can in itself impact population persistence (Lande et al., 2003; Postuma et al., 2020). Teasing out the impact of explicitly considering transgenerational effects for population dynamics therefore requires the careful comparison of multiple models. To explore the role of transgenerational effects within a tightly-coupled trophic system responding to generation-scale variance in temperature, we first conducted highly replicated experimental trials to assess how the thermal performance curves of both host and parasitoid are influenced by the temperatures the previous generation is exposed to. We test whether including the temperature experienced by the previous generation can improve predictions of reproductive rate. Focussing on the impacts on the population persistence under warming conditions, we then used the observed responses to parametrise simulation models to explore the impact of the observed transgenerational effects.. We hypothesised that there will be an extra emergent dynamic impact of the combined transgenerational effects across trophic levels.

## Methods

### Study organisms

Drosophila (Diptera: Drosophilidae) provide a practical system that has been widely used to study both thermal performance and community responses to climate change (Davis et al., 1998; Hoffmann & Sgrò, 2011; Rezende et al., 2020). In *D. melanogaster*, a significant interaction between parental and offspring thermal environment on the entire thermal performance curve has been previously identified (Cavieres et al., 2019; Gilchrist & Huey, 2001; Huey et al., 1995). Compared to other model systems of phenotypic plasticity (e.g. Fey et al., 2021) our use of *Drosophila* allows for the discretisation of generations and a simplification of the modelling process.

We used *Drosophila sulfurigaster* (Duda, 1923) and a generalist hymenopteran parasitoid wasp *Asobara* (Braconidae: Alysiinae, strain: KHB, reference: USNMENT01557097, BOLD process ID:DROP043-21, species identifier: drop_Aso_sp8, Lue et al., 2021). *D. sulfurigaster* is a partially susceptible native host of *Asobara* parasitoids, allowing for some variation in the rate of successful parasitism across treatments. The parasitoid *Asobara*, attack the second-instar larval stage, with a single offspring emerging from each host pupa. Importantly *Asobara* (henceforth simply ‘wasps’) have approximately the same generation time as *D. sulfurigaster* (henceforth simply ‘flies’) simplifying both experiments and analysis. Prior to the experiment, the wasps had been maintained on highly susceptible *D. melanogaster* larvae.

Both host and parasitoid species were established from adults collected in the Wet Tropics bioregion of northeast Queensland, Australia. There, the *Drosophila* community suffers considerable parasitoid pressure with wasp DNA found in between a third to a half of pupae (Jeffs et al., 2021). *Drosophila* and parasitoid cultures were originally exported under permit PWS2016-AU-002018 granted by the Department of Energy and Environment of the Australian Government. These cultures had been maintained at a constant 24°C at the Biology Centre, Czech Academy of Sciences, before being transferred to the Department of Biology, University of Oxford, where they were maintained at 25°C on cornflour–sugar-yeast–agar *Drosophila* food medium in a 12:12h light: dark cycle for the preceding year.

### Effect of temperature on fly reproduction

The preceding generation of flies (G0) was initiated in 48 bottles each with 30ml of food medium. Approximately 20 stock adult flies were allowed to lay eggs for 24 hours to generated a low density in each bottle. These bottles were then randomly assigned to incubators set at one of three temperatures (19, 23, 27°C), under a 12:12h light:dark cycle (Figure 1). This 8°C range was chosen based on prior knowledge of the thermal sensitivity of this community (Chen & Lewis, 2024) to be ecologically relevant and approach the reproductive limit of the host. Mean temperatures at the focal area both species originate from range from 21°C at the highest elevation sites to 26°C at the lowest (Jeffs et al., 2021). Temperature and humidity loggers in each incubator showed that a controlled environment was maintained throughout the experiments. Across all the experiments, pupation cards were added to provide a vertical surface for pupae.

**Figure 1.**
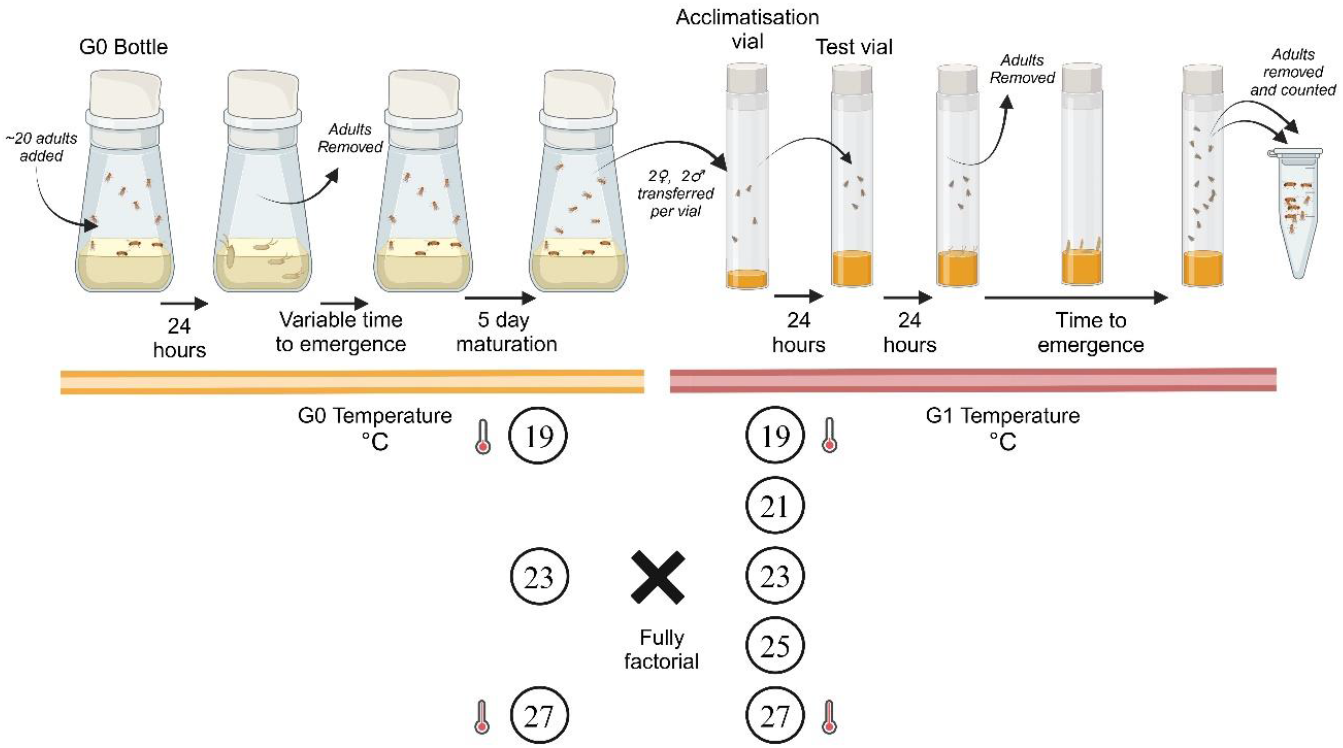
Summary of experimental design for the fly temperature treatment experiment. G0 = Generation 0. G1 = Generation 1. Large numbers of G0 bottles were initiated with approximately 20 flies and randomly assigned to incubators set at one of three G0 temperatures (19, 23, 27°C). After 24 hours to lay eggs, the adult flies were removed. Once G0 flies emerged and matured, two males and two females were counted into each acclimatisation vial. Acclimatisation vials were randomly assigned to incubators set at one of five temperatures (19, 21, 23, 25, 27°C) in a fully factorial design (n=40 per combination). After 24 hours of acclimatisation, flies were transferred into standardised test vials that remained in their assigned G1 test temperature to lay eggs for 24 hours. G1 emergences were regularly removed and counted.

Subsequent steps were timed relative to developmental milestones, as time to eclosion is dependent on the temperature treatment (nine days at 27°C, 11 days at 23°C and 15 days at 19°C). Since *D. sulfurigaster* can take four days post-eclosion to mature enough for egg laying (J. Chen, *pers. obs.*), a maturation time of five-days post-eclosion was allowed in the G0 temperature before two male and two female G0 flies were counted into acclimatisation vials containing food medium under light CO_2_ anaesthesia. Forty such acclimatisation vials from each G0 temperature were randomly assigned to incubators set at one of five temperatures (offspring generation (G1) temperatures: 19, 21, 23, 25, 27°C) for 24 hours (600 vials total). The acclimatisation vials minimised the impact of the CO_2_ anaesthesia and allowed acclimation to the new temperature.

For the reproduction trials, the flies in each acclimatisation vial were transferred into standardised test vials containing 5ml of food medium. Any deaths at this stage were noted, but otherwise considered part of the performance assessment. These vials remained at their G1 test temperature. After 24 hours of egg laying, the G0 adult flies were removed. Vials were checked regularly, and as G1 flies emerged, they were removed, stored, then counted. In total, 15,755 flies were counted across the 600 trials.

### Effect of temperature on wasp reproductive output

The wasp temperature treatment experiment followed a similar design (Figure 2). Standardised host vials were founded with five adult female flies, sufficient males, 5ml of standard food medium and a sprinkle of yeast granules to stimulate egg laying and reduce variance in egg laying rate. Vials were maintained at 25°C and adult flies were allowed to lay eggs for 48 hours before being removed. Two male and two female wasps were aspirated into each standardised host vial shortly after initiation to attack the early instar larvae. Based on trials with fully susceptible hosts we expected this ratio to result in the great majority of hosts being attacked by the wasps. A total of 48 G0 host vials with wasps were randomly assigned to one of the three G0 temperatures (19, 23, 27°C, 16 per temperature).

**Figure 2.**
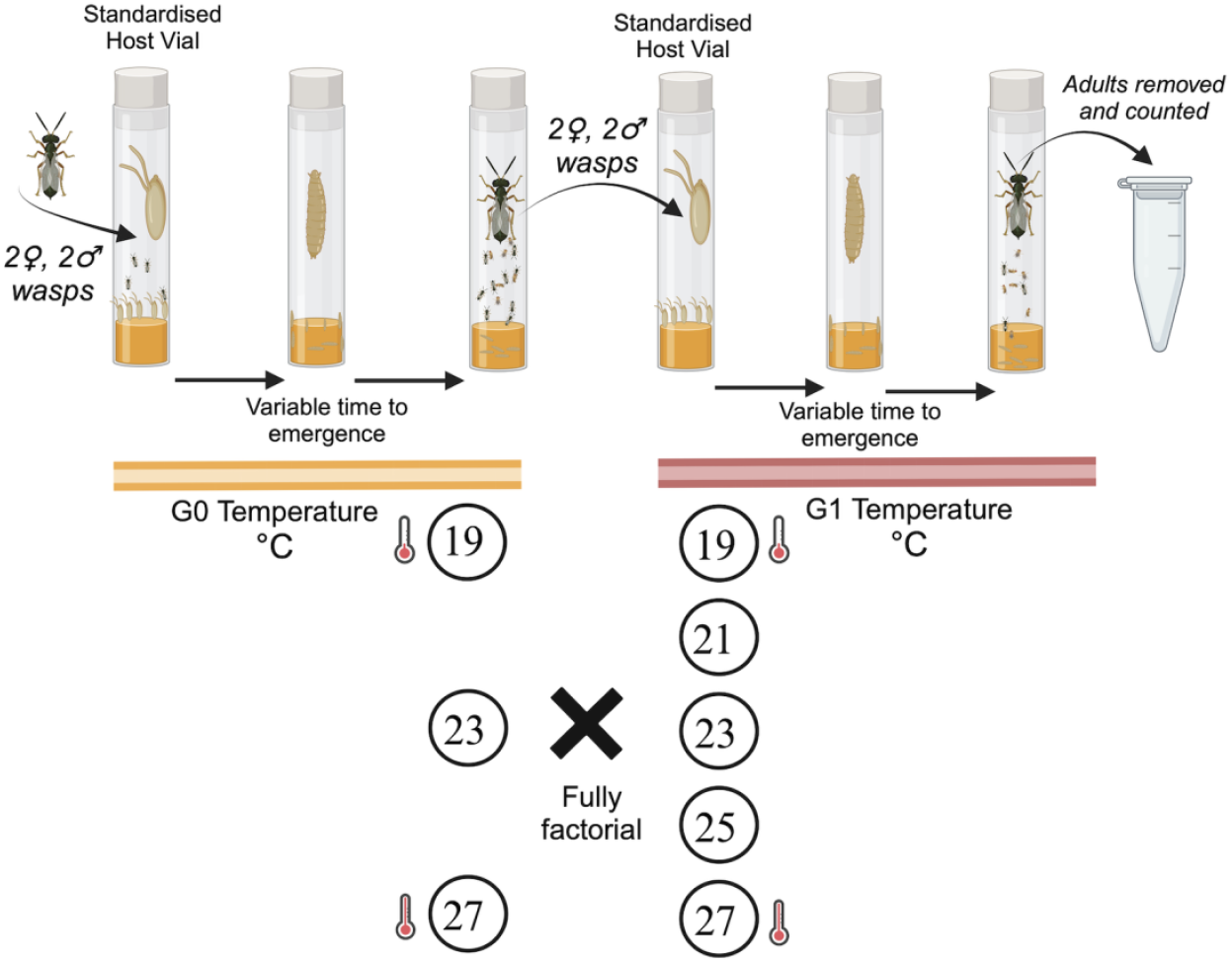
Summary of experimental design for wasp temperature treatment experiment. G0 = Generation 0. G1 = Generation 1. To initiate G0, two male and two female wasps were added to standardised vials of hosts. The vials were stored in incubators at either 19, 23, 27°C. Two male and two female wasps recently emerged were transferred to G1 host vials randomly assigned to G1 temperatures (19, 21, 23, 25, 27°C), in a full factorial design. Each treatment combination had between 16 and 20 replicates. As G1 wasps and flies emerged they were removed, stored and then counted.

Due to differences in development time, the timing of G1 was staggered. Upon the first observed adult wasp emergence from each G0 treatment, G1 standardised host vials were prepared (as described above), to which two male and two female G0 wasps were added. These vials were randomly assigned to one of the five G1 test temperatures (19, 21, 23, 25, 27°C), following the fully factorial design. There were sufficient G0 wasps for 16-20 replicates for each treatment combination (totalling 277 vials across all 15 combinations). Test vials were checked regularly and both flies and wasps removed, stored and counted. In total, 6564 flies and 18,453 wasps were counted.

### Statistical analysis

All statistical analyses were performed in R version 4.4.0 (R Core Team, 2024).

#### Flies

To test the statistical support for parental generation (G0) temperature influencing fly reproduction rate in the subsequent generation (G1), multiple models were fit and compared by AIC, with and without the inclusion of G0 temperature. A thermal performance curve likelihood function (Equation 1) based on the deutsch_2008 function (Deutsch et al., 2008) in the rTPC package (Padfield, 2023) was customised to fit to discrete count data, and fit using the bbmle package (Bolker et al., 2023).

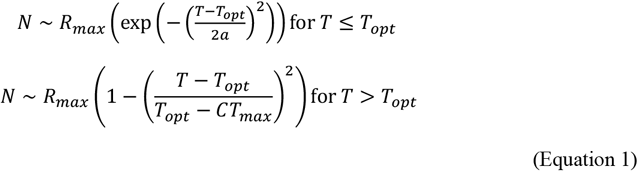

where *N* is the number of flies expected and *T* is the G1 temperature (°C). This function reproduces the expected shape of the thermal performance curve with biologically interpretable parameters. The three fitted parameters are: the optimum temperature *T_opt_*, the maximum reproduction at optimum temperature *R_max_*, a curve width parameter *a*. The critical thermal maximum *CT_max_*fell outside the range of the experimental trials and was fixed at 30°C based on previous studies in this species (Chen & Lewis, 2024). To account for the overdispersion in the observations, a negative binomial error function was used.

The model variants were tested where the three fitted parameters (*T_opt_, R_max_, a*) were fixed regardless of the G0 temperature (Model 1b), with the parameters having a linear dependence on G0 temperature (*θ*(*T*) = *θ*_19_ + (*T*_*G0*_− 19)*β_θ_*, Model 1c), and with the parameters having separate values for each of the three G0 temperature levels (Model 1d). Three further models were tested that included temperature dependence on only two of the three parameters (Models 1e-g).

#### Wasps

A similar approach was followed to test if G0 temperature influences wasp reproductive output. To account for variation in the numbers of larvae in the standardised host vials, the proportion of wasps of the total emerged insects (wasp count/ (wasp count + fly count)) was used, assuming a binomial distribution. This approach assumes that the combined count was independent of G1 temperature, which was supported (linear model of Total Count ∼ G1, *F*_1,275_ = 0.373, p = 0.541).

Unlike for fly reproduction, there is no established thermal performance model to use for parasitoid wasps. Based on inspection of the data, a monotonic relationship between temperature and reproduction appeared justified within the trial range and binomial generalised linear models were fit. Multiple models were fit and compared by AIC: without inclusion of G0 temperature (Model 2a), with G0 temperature as a linear predictor (Model 2b), with an interaction between G0 and G1 temperature (Model 2c). To examine if the effect of G0 was non-linear, separate models for each of the three G0 temperatures were fit and their likelihoods combined (Model 2d).

### Simulation Modelling

To explore how transgenerational effects of thermal conditions and autocorrelation in temperature may impact the time to extinction of this host-parasitoid system under a high-emissions climate change scenario, we built simulation models incorporating data from the experiments. Both host and parasitoid species are assumed to have discrete and synchronised generations in a Nicholson-Bailey framework (Hassell, 2000).

Separate simulation models were built parameterised either with the model fits with or without including transgenerational effects in either species’ performance. Two modelling frameworks were used to explore the potential impact on dynamics, one very simple using Type II functional response, the second using a somewhat more realistic representation of spatial aggregation.

#### Model A – Type II Functional Response

The potential fly population in the next time step 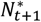, given the current fly population density *N_t_* is based on a logistic growth model:

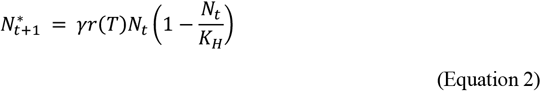

where potential fly population growth rate in a particular environment *r*(*T*) is scaled by a failure rate *γ,* and *K_H_*is a host carrying capacity term.

The number of flies lost to parasitism followed a Type II functional response:

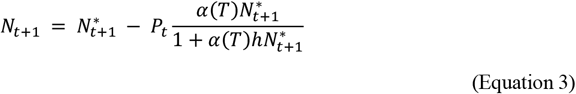

where *h* is a handling time, and *α*(*T*) is the temperature dependent attack rate.

The wasp population in the next generation was calculated similarly, with the addition of a small immigration component (*d*) to maintain some wasp threat throughout:

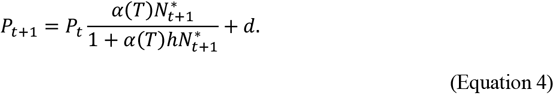

#### Model B – Proportion surviving under Spatial Aggregation

An alternative model followed the overall structure of Model A, but models the conversion of potential hosts as a proportion that survive assuming aggregated parasitoid attacks determined by the parameter *k*, following Hassell (2000) and Meisner et al. (2014):

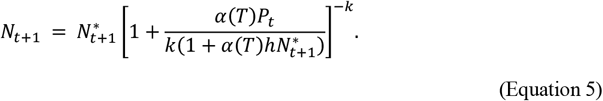

Taking the wasp population as those that do not survive, the wasp population size in the next generation is then:

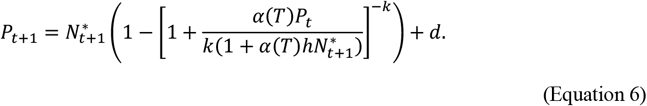

#### Experimentally Derived Temperature Dependence

The fly population growth rate *r*(*T*) followed the environmental performance curve fit to the empirical data (Equation 1). Without the inclusion of transgenerational effects, the values of the thermal performance curve parameters were fixed to those identified with Model 1b (Table S3). When transgenerational effects are included, linear interpolation is used between and beyond the curve parameter values identified by the best fit model (Model 1d, Figure S1). To prevent negative *R_max_* values, the expression was lower-bounded at 0.1.

To represent temperature dependence in the attack rate *α*(T) of the wasps, the proportion models fit to the experimental data (*η*(*T*)) were scaled by a maximum wasp attack coefficient *t_p_*:

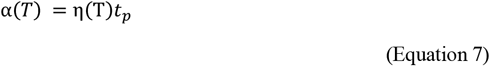

Model 2a (which did not include G0 temperature as a predictor) was used for *η(T)* without the inclusion of transgenerational effects. To include transgenerational effects, a model equivalent to Model 2d was used which interpolated between the fitted values of the parameters at the three G0 with a quadratic function.

#### Simulation model parameter calibration

For both models, remaining parameters were selected based on a combination of experimentally derived values, previous literature and manual tuning to generate plausible dynamics prior to the introduction of simulated climate change (Table S3). Fly carrying capacity *K_n_*was set sufficiently large to overcome demographic stochasticity, whilst fly failure rate *γ* was set on the same order of magnitude as *R_max_*to stabilise the overall growth rate and prevent overly-explosive exponential growth. Wasp handling time *h* was set at 0.01 as, on average, the wasps can parasitise a maximum of approximately 100 flies – due to egg limitation (maximum attack rate is 1/*h*) (Godfray, 1994; Hassell, 2000). The wasp attack coefficient *t_p_*is a scaling factor that defines wasp attack rate and was tuned to generate an initial rate of parasitism of approximately 50% in line with field studies of this system (Jeffs et al., 2021). The wasp spatial aggregation term *k* was set last, with simulations run at a range of values around that in Meisner et al. (2014) to generate plausible dynamics with the populations varying moderately around their carrying capacities.

#### Generating simulation temperature regime

Simulated temperature trajectories were built on a 500 generation burn-in period with mean temperature 24°C (to settle any initial transient dynamics), followed by a linear rise of 5°C over 2000 generations (approximately 80 years). On top of this, Gaussian noise (σ =1) was superimposed, with an autocorrelation of either 0 or +0.8 (Figure 5). To prevent numerical errors outside the bounds of the extrapolation, temperatures were capped at 30°C, the flies thermal maximum.

These values were selected to align with the ranges and coefficient of variation observed in fortnightly averages from the summer months at a long-term meteorological station near the collection site of the study species, Koombooloomba Dam (Australian Government Bureau of Meteorology, 2023) and a high emissions warming scenario (IPCC, 2023). The extended temperature ramp minimises lag-effects and therefore also allows our results to be approximately interpreted as the mean temperature at which the community is able to persist at.

#### Simulation trials

The simulation models were used to investigate the time to extinction of the hosts under a suite of model assumptions: the inclusion of transgenerational effects in each of the flies and wasps, and the inclusion of positive autocorrelation in temperature between generations. Each model scenario was run 100 times with consistent sets of random seeds to generate the temperature trajectories used across scenarios. Extinction was defined as the first generation in which the fly population fell below one.

To test if there was an interaction between the impact of transgenerational effects (in the flies and wasps) and autocorrelation in temperature on (logged) time to extinction, a linear mixed effects model was fit using the *lme4* package (Bates et al., 2015). By taking the logarithm, we modelled the effect of the treatments on the proportional change in time to extinction, thus assuming a multiplicative relationship between variables. The full model included as fixed effects autocorrelation (0 or 0.8) and the inclusion of transgenerational effects for each species (ON/OFF), along with all interaction terms. Temperature trajectory seed ID was included as a random effect. The significance of the interaction terms was tested with a Type III ANOVA with Satterthwaite’s method via the *lmerTest* package (Kuznetsova et al., 2017).

## Results

### Experimental Evidence

For the fly temperature treatment experiment, the best model fit included G0 temperature as an explanatory variable with all parameters able to vary with the three levels of G0 (Table 1, Model 1d), indicating a significant impact of transgenerational effects of thermal conditions on fly performance. As G1 temperature increased, emerged offspring numbers increased for all three fly G0but at distinct rates (Figure 3). At a G1 temperature of 19°C, the greatest predicted reproduction was seen when G0=19°C. At all temperatures above 19°C, the greatest predicted reproduction was seen when G0=23°C. When G0=27°C, performance was low across the temperature range, suggesting the presence of a non-adaptive condition-transfer effect.

**Table 1.**
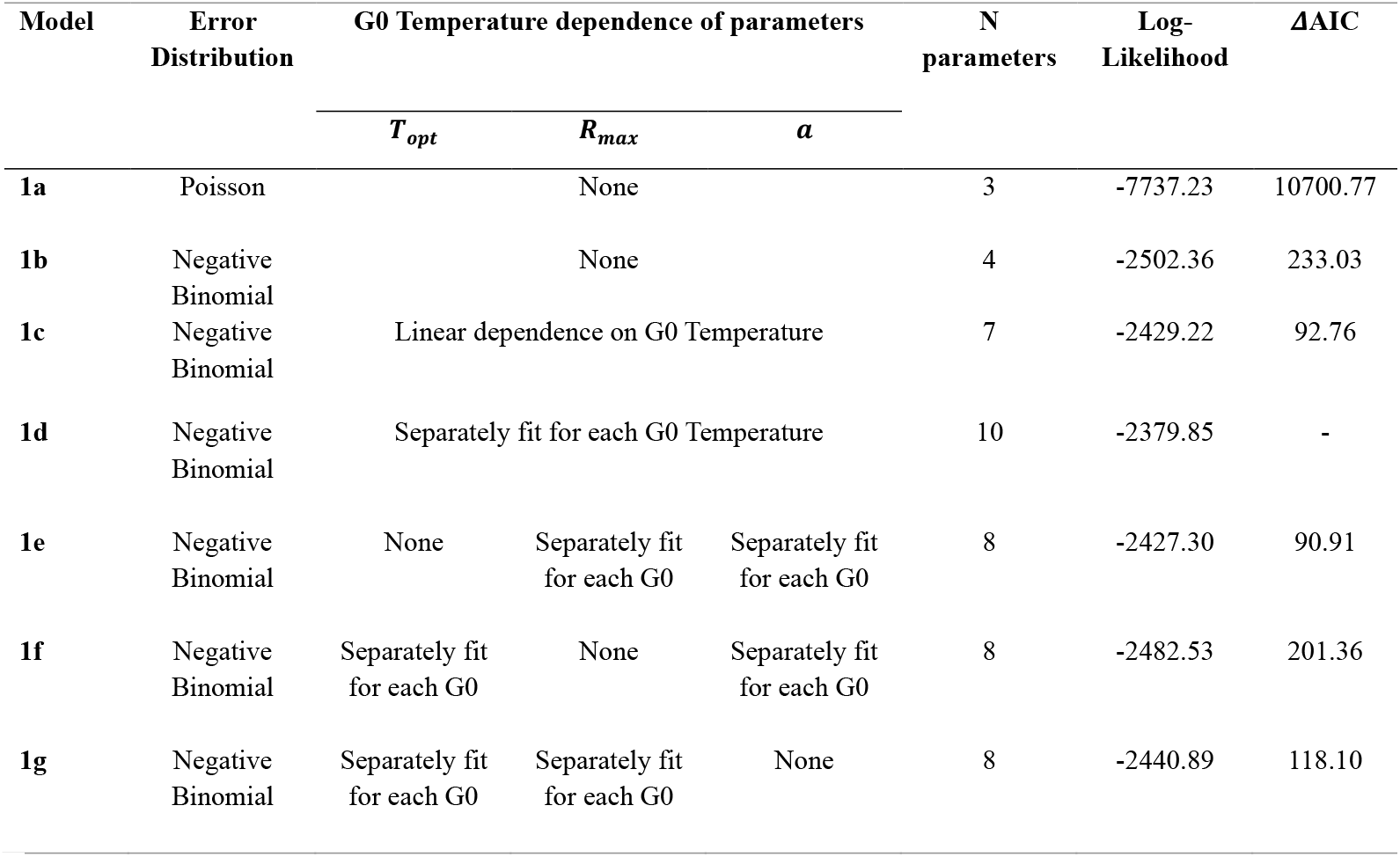
Tests of impact of G0 temperature on fly reproduction rates through model comparison. The best supported model (1d), included full non-linear dependence of the three fitted thermal performance curve parameters on the temperature in the previous generation (G0). Models that included the G0-temnperature dependence for only two of the three parameters (Models e-g), or included a linear G0 temperature dependence (1e) gave notably less good fits (ΔAIC > 90). Using a simpler Poisson error distribution (1a) was not supported at all.

**Figure 3.**
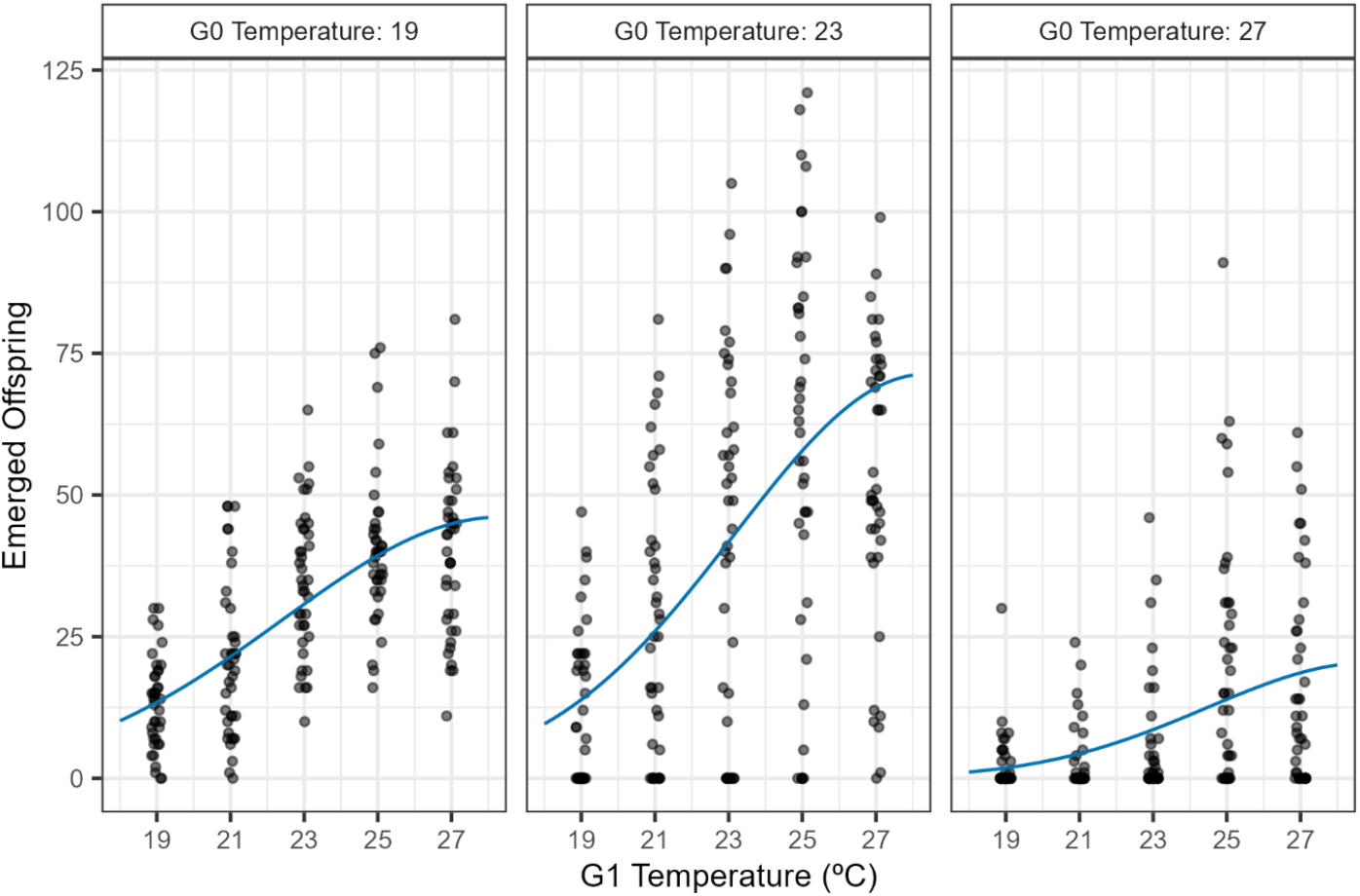
Fly Reproduction Best Fit Thermal Performance Curves and Raw Data. Each panel shows the effect of temperature during that generation (G1) on the reproductive rate (number of emerged offspring) for each temperature in the preceding generation (G0). See Figure 1 for how generations were delineated. Black points represent raw data from each vial and have been jittered horizontally. Means and standard errors for each treatment combination are given in Supplementary Figure S3. Blue lines represent central expectations based on the best fit thermal performance curve model. Parameter estimates for the thermal performance curves and 95% confidence intervals are in the supplementary information (Table S1).

For the wasp temperature treatment experiment, the best supported model included G0 temperature as a categorical factor with three levels interacting with the effect of G1 temperature, indicating a significant transgenerational impact on wasp performance (Table 2, Model 2d). In comparison to the flies, the association between G1 temperature and wasp performance was more variable depending on G0 temperature (Figure 4). When G0=19°C, as G1 temperature increased, the predicted proportion of wasps decreased. In contrast, when G0=27°C, as G1 temperature increased, the predicted proportion of wasps increased. This could be interpreted as an adaptive transgenerational effect.

**Table 2.** Tests of the impact of G0 temperature on wasp performance through model comparison. **The best model** (2d), included distinct responses to the temperature in the previous generation (G0) and was much more strongly supported than models that included G0 temperature as a linear predictor. G0 = Temperature of wasp generation 0. G1 = Temperature of wasp generation 1.

| Model | Explanatory variable(s) | n | Likelihood | ΔAIC |
| --- | --- | --- | --- | --- |
| 2a | G1 | 2 | -4913.71 | 1364.77 |
| 2b | G1+G0 | 3 | -4552.96 | 645.28 |
| 2c | G1*G0 | 4 | -4397.26 | 335.87 |
| 2d | G1*factor(G0) | 6 | -4227.32 | - |

**Figure 4.**
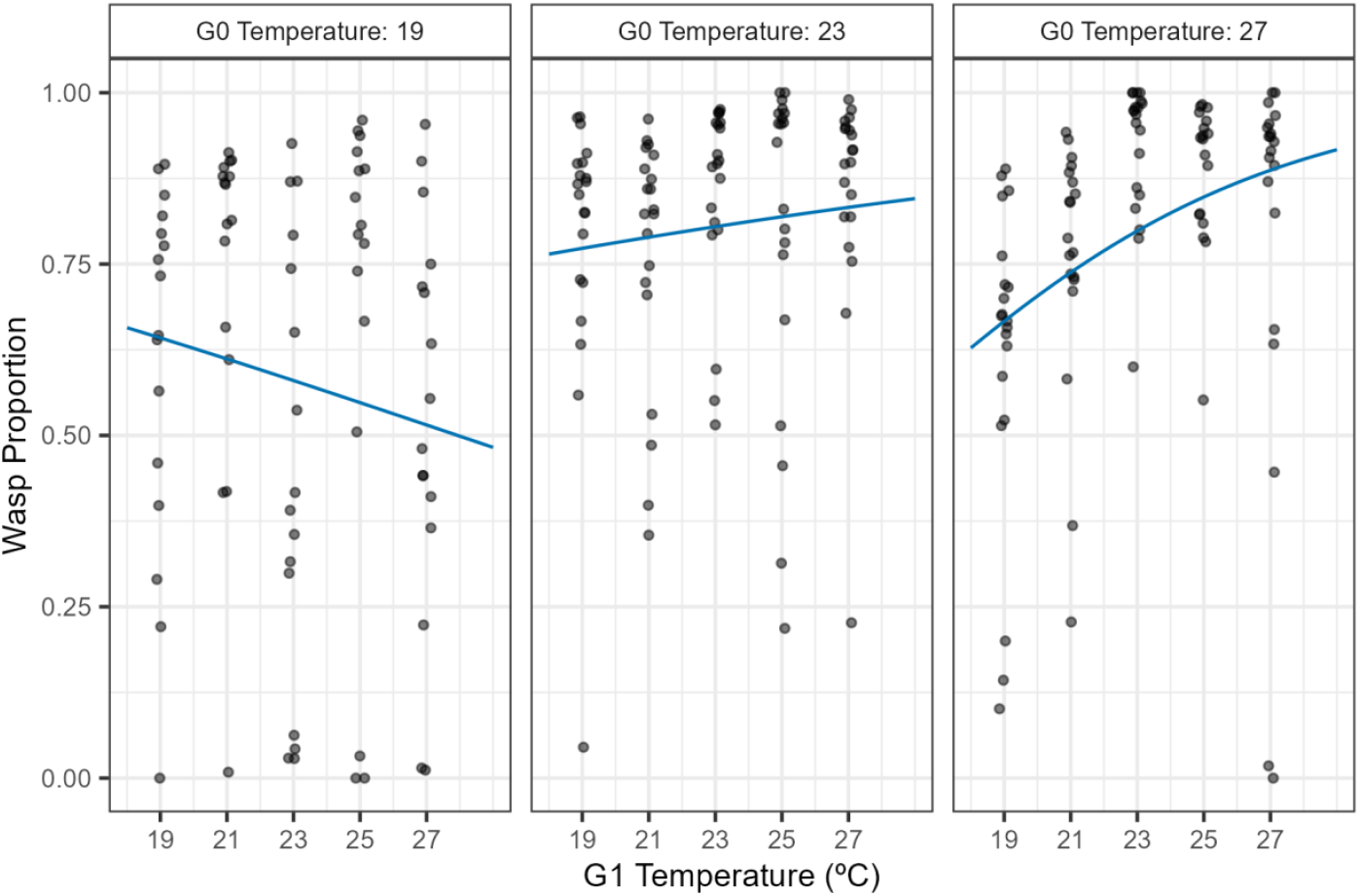
Wasp Emergence Rate Best fit Thermal Performance Curves and Raw Proportion Data. Each panel shows the effect of temperature on the proportion of wasps (wasp count/total count, where total count includes the flies and G1 wasps) for each previous generation temperature (G0). See Figure 2 for delineation of generations. Black points represent raw data from each vial and have been jittered horizontally. Means across the vials and approximations of standard errors for each treatment combination are given in Supplementary Figure S4. Blue lines represent predictions based on best fit model. Fitted model parameter estimates and 95% confidence intervals are in the supplementary information (Table S2).

### Simulation model results

Both autocorrelation and the full inclusion of transgenerational effects markedly hastened the extinction of the fly host in the simulations under both models (Figure 5, Table S4). Converting from generations to times, accounting for transgenerational effects hastened the time-to-extinction by approximately 30 years and reduced the temperature at which they could persist at by nearly 2°C. In the simpler Model A, autocorrelation and each of the transgenerational effects (i.e. the fly and the wasp) had a significant interactive effect (three-way interaction term, *F*_1,693_= 10.86, p = 0.001, Table S5). Taken separately, transgenerational effects on the fly had a markedly greater impact. Transgenerational effects on the wasp alone had minimal impact, but in combination with transgenerational effects on the fly and autocorrelation, further hastened extinction (Figure 5, top left). However, under Model B there were no detectable interactive effects between either of the transgenerational effects and autocorrelation on the log-time-to-extinction (*F*_1,693_= 0.0083, p = 0.927, Table S5).

**Figure 5.**
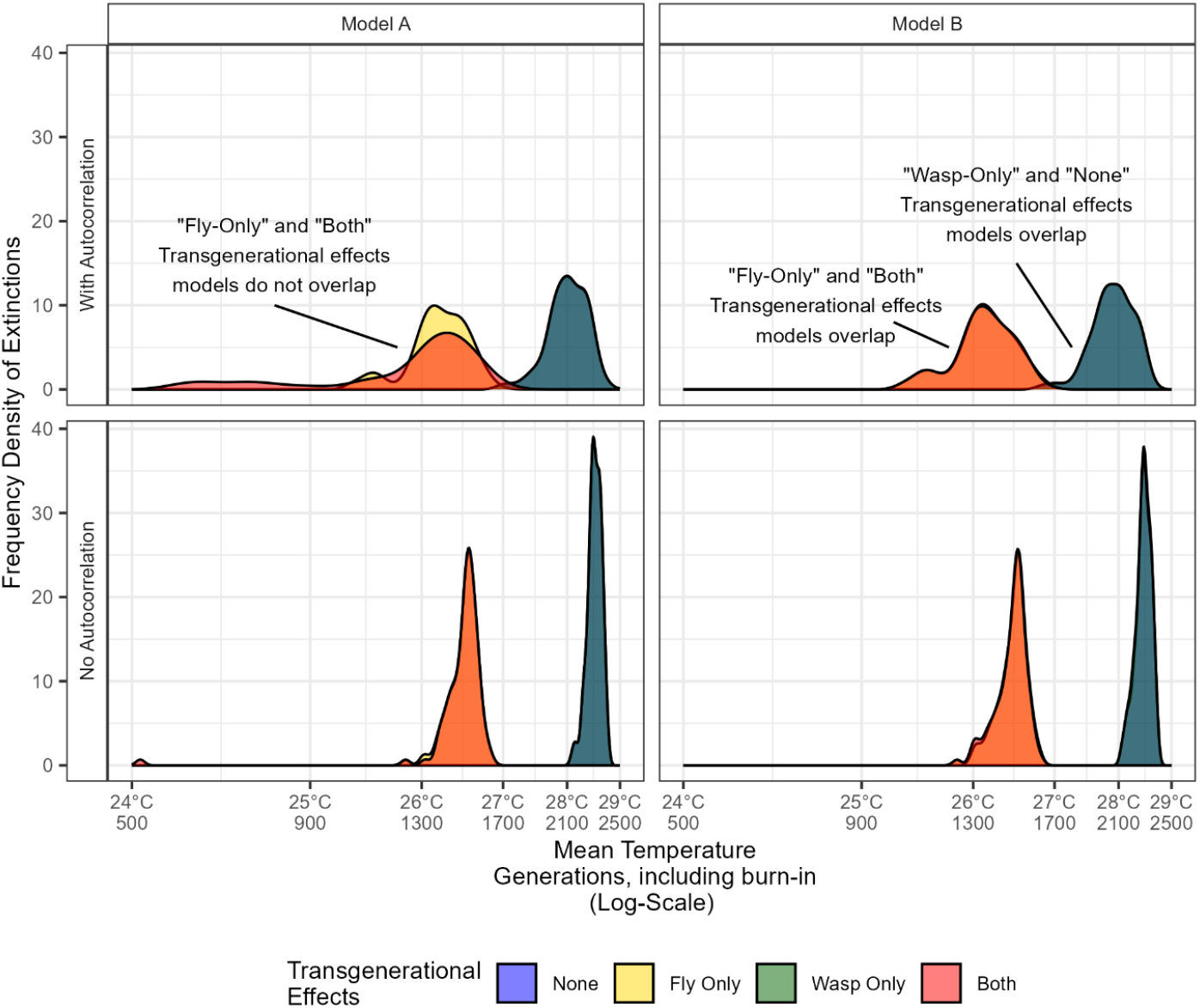
Identification of dynamic effect of combined transgenerational effects emerging from joint inclusion of environmental autocorrelation and both identified transgenerational effects. Density histograms of the observed distribution of time to extinction (in generations), over 100 simulations the host-parasitoid system under different scenarios and models. There is a near perfect overlap between scenarios with transgenerational effect modelled for ‘wasp-only’ and ‘none’ (blue and green, combining to dark turquoise), as well as ‘both’ and ‘fly-only’ (yellow and red, combining to orange), with the exception of the top left panel that uses a simpler functional response and includes autocorrelation. Mean temperature increases by 1°C per 400 generations. Geometric means are given in Table S4. Note the logarithmic x-scale.

## Discussion

These results demonstrate how transgenerational effects can have substantial impacts on the consequences of climate change through a variety of pathways. Despite hope and interest in the potentially resilience-enhancing nature of adaptive responses (Sohlström et al., 2021), our results highlight the negative aspects of transgenerational effects. Modelling is crucial to contextualise the various processes (Terblanche & Hoffmann, 2020), but our observation of only seeing emergent dynamic consequences of transgenerational effects with certain model forms emphasises the need for caution and further theoretical developments.

Two key caveats need to be reiterated when interpreting these results: firstly, our focus was on identifying potential impacts of cross-generational transfer of population dynamics rather than isolating physiological mechanisms. As such, although the nature of these responses suggests contributions from both condition-transfer effects and potential adaptive transgenerational plasticity, these cannot be cleanly differentiated within our design. This is a fundamental challenge – while the observed effects are ‘transgenerational’ from the perspective of standard population models that track adults (Hassell, 2000), it is plausible that the most important underlying physiological processes can be considered within-generational (Donelson et al., 2018) from other perspectives or framings (e.g. egg-to-egg). Secondly, these models are not intended to be specific representations of particular real-world scenarios or locations. The experiments focus on just two species from a complex community (Chen & Lewis, 2023, 2024; Jeffs et al., 2021) and do not account for many features that would affect climate change responses against a fluctuating environment, from microclimates (Pincebourde & Woods, 2020), competition (Terry, 2025), the impact of seasonality (Tougeron et al., 2020), to the potential for evolution (Chevin & Bridle, 2025; Rudman et al., 2022).

### Experimental observations

We found clear experimental evidence that the parental thermal environment had a significant and substantial impact on subsequent reproductive output in both the hosts and parasitoids. The somewhat discordant responses of the host reproductive output and the parasitism rates to temperatures across generations opens up the potential for complex dynamics to emerge.

When the parental fly generation was maintained in a thermally stressful environment (G0=27°C), subsequent reproductive output was lower across the temperature range (Figure 3), in line with previous results using *D. melanogaster* that identified an elevated cost to living in an extreme thermal environment, with high and constant temperatures reducing the thermal performance of the offspring generation (Cavieres et al., 2019). The most direct explanation is that G0 flies that developed in stressful temperatures were not able to take advantage of more clement conditions in the G1 generation. On average, across all offspring generation temperatures, the best fly performance was when the parental generation was maintained at an intermediate temperature (G0=23°C, Figure 3). This indicates a condition-transfer effect and aligns with the optimum parental temperature hypothesis that parents kept at intermediate temperatures produce fitter offspring compared to parents at temperature extremes (Cohet & David, 1978; Gilchrist & Huey, 2001). Although our experimental design cannot distinguish developmental success rates from the number of eggs laid, it does reiterate the importance of accounting for potential carry-over effects when assessing thermal performance (Jenkins & Hoffmann, 1994; Schiffer et al., 2013; Sgrò and & Hoffmann, 1998).

There was also some indication of adaptive cross-generational effects where offspring facing the same environmental conditions as their parents have a higher fitness than offspring facing different environmental conditions (Gilchrist & Huey, 2001) in anticipation of what they are likely to experience (Cavieres et al., 2020; Tariel et al., 2020). For instance, when the parental generation was maintained in a thermally stressful environment (G0=27°C), the proportional decrease in reproduction was lower when the offspring generation was also in a thermally stressful environment (Figure 3). This could indicate the possibility of some anticipatory effect, although it is overwhelmed by the non-adaptive transfer of poor condition. Furthermore, the greatest thermal performance at 19°C was when the parental generation was also maintained at 19°C (Figure 3). This highlights that while it is important to differentiate between anticipatory and condition-transfer effects when understanding the evolutionary causes and consequences of transgenerational effects (Sánchez‐Tójar et al., 2020), both can operate simultaneously.

The assay we used (with a discrete window of egg laying time) is especially sensitive to the rate of egg laying but does not capture longevity of the G0 flies. While fly reproduction was lower when the parental generation was maintained in a thermally stressful environment, demographic trade-offs may have meant other traits, such as thermal tolerance (Bozinovic et al., 2011) or longevity (Ismaeil et al., 2013), were not. In a previous study, *D. melanogaster* subjected to a variable and stressful thermal environment exhibited reduced reproduction, yet showed increased thermal tolerance compared to flies kept at a constant temperature (Cavieres et al., 2020).

The wasp thermal performance curves show clear evidence for improvements in thermal performance where there was a close temperature match between the generations. When the parental generation was maintained at a low temperature (G0=19°C), the greatest wasp proportion was when the G1 temperature was also low (Figure 4). In comparison, when the parental generation was maintained at a high temperature (G0=27°C), the greatest wasp proportion was when the temperature was also high. This acclimation would be expected to reduce their vulnerability to climate change despite many of their life-history traits, such as a high trophic position and typical trophic specialisation, increasing their susceptibility (Jeffs & Lewis, 2013). We cannot however concretely identify if these changes are due to changes in host resistance, wasp virulence, or changes to the functional response with our experimental design. We also note that in our experimental design using standardised host vials, we were not able to test if there was an interactive effect between the parental temperature of the host and their susceptibility to the wasp.

### Time to extinction: Role of transgenerational effects and autocorrelation in temperature trajectories

The simulation models predict that including transgenerational effects of thermal conditions significantly hastens the time to extinction of this host-parasitoid system. The substantial negative effect of including the transgenerational effects in the simulation models highlights the predominance of negative condition-transfer over any potential adaptive transgenerational plasticity through anticipatory effects. Autocorrelation, even without transgenerational effects, magnifies the threat posed by environmental stochasticity (Lande et al., 2003). By prolonging periods of adverse conditions and reducing the presence of temporal refugia, positive temporal autocorrelation can amplify environmental stress and exacerbate the risk of extinction (Postuma et al., 2020).

Without autocorrelation, there *should* be no special dynamic consequences of transgenerational effects. The observed strong impact seen in the scenarios without autocorrelation show that effectively averaging across assays of the thermal performance of flies raised in more suitable environment is inflating the estimated performance at higher temperatures. The additional effects from the combination of autocorrelation and transgenerational effects are the direct dynamic impact from the transgenerational effects. In the simpler Model A, there was a clearly identifiable additional interaction effect, although this dynamic component was considerably smaller than the independent impacts of both autocorrelation and the direct effects of transgenerational effects. The modelled transgenerational effects observed in the wasps had essentially no impact on the time of fly extinction on their own, but under Model A did have an impact in combination with the transgenerational effects observed in the flies. The double-effect of sequential host generations of reduced host and improved wasp performance brought forward the point at which the host population was pushed below the extinction threshold. This negative effect highlights how any benefits of acclimation were overwhelmed by condition-transfer effects. By contrast, no interactive effect is detected within Model B. Both models are relatively standard representations of host-parasitoid interactions, while Model B assumed spatial clumping through the negative binomial term, which might act as one of the buffering mechanisms that absorb the emergent effect. The distinct responses under the two models highlights the context dependency of whether experimentally identified effects will have population-level consequences.

### Linking experiments, physiological mechanisms and dynamic consequences

Precisely distinguishing the multiple possible contributing factors that can underly acclimatory and transgenerational effects is a significant challenge (Terblanche & Hoffmann, 2020), further confounded by difficulties precisely delimitating the boundaries between generations (Burggren, 2015; Lee et al., 2020) and the potential for influences to span multiple generations (Tariel et al., 2020). For example, environmental impacts on primordial germ cells that appear as examples of transgenerational plasticity could be due to within-generation plasticity in offspring during early stages of development (Donelson et al., 2018).

Our broad-scale approach allows us to deploy the high replication necessary to confidently identify effects but cannot differentiate physiological mechanisms, and the experimental requirements to fully partition all potential different processes can quicky become unfeasible at scale. While further experiments could address some of the gaps, for example transferring a controlled number of eggs to differentiate the impact of adult condition influencing egg laying rate from effects related to offspring development at different temperatures, it is hard to assess how much difference such extra information would bring. As such, our results should be taken as experimentally-informed potential scenarios, rather than precise forecasts. In scenarios with stochastic environments and multiple interacting species, model predicted outcomes are readily qualitatively changed under different parameter combinations (Terry et al., 2022). Closing the complexity gap and determining the appropriate level of abstraction to tackle these processes at scale will be a key challenge for future research. Our experiments were designed to align with models of host-parasitoid dynamics that conventionally operate on an adult-adult basis (Nicholson & Bailey, 1935), but demarking the generational boundary at a different stage (e.g. eggs – eggs) could well give different results. Insect life stages are known to be impacted differently by temperature (Kingsolver et al., 2011; Pawar et al., 2024) and so our generational scale thermal treatment could be missing important consequences by not accounting for demographic structuring (Amarasekare & Coutinho, 2013).

## Conclusion

Our results empirically highlight the substantial, but context-dependent, impact multi-trophic transgenerational effects can have on population dynamics under climate change. These findings underscore the need to quantify and contextualise transgenerational effects when understanding species’ responses to climate change and show the need to both move beyond demonstrations that such effects exist to improving our theoretical understanding of the scenarios in which they are influential.

## Acknowledgements

We thank the support of the CERO and the Fly Lab groups, the Hrček lab for sharing lines, as well as comments and advice from Owen Lewis. JCDT was funded by the Leverhulme Trust ECF-2022-666. JC was funded by NERC NE/X000117/1. Figures 1 and 2 created with BioRender.com.

## Author Contributions

NLB conducted the experiments and wrote the first draft of the manuscript. JCDT supervised the project, designed the analyses and finalised the manuscript. JC co-designed the experiment and significantly contributed to the finalisation of the manuscript. All authors contributed critically to manuscript development and gave final approval for publication.

## Data Availability

Raw data, code, fitted model objects and markdown documents covering all steps are publicly archived on Zenodo https://doi.org/10.5281/zenodo.13327764 and available through GitHub at https://github.com/jcdterry/MultiGenHeatEffects_public.

