## Supplementary material for "Transgenerational effects increase the vulnerability of a host-parasitoid system to rising temperatures": SI Figures and Tables

**Table S1: Parameter estimates and their 95% confidence intervals for the fly thermal performance curves.** * Denotes that the 95% confidence interval reached the upper boundary set in the model. For $T_{opt}$ this upper boundary was set at 29°C due to $CT_{max}$ being at 30°C.

| Parameter | Estimate | 2.5% | 97.5% |
|---|---|---|---|
| **$T_{opt}$19** | 28.37 | 24.38 | 29.00* |
| **$T_{opt}$23** | 28.35 | 24.07 | 28.99 |
| **$T_{opt}$27** | 29.00 | 26.81 | 29.00* |
| **$R_{max}$19** | 46.11 | 34.76 | 58.44 |
| **$R_{max}$23** | 71.31 | 48.67 | 76.15 |
| **$R_{max}$27** | 20.50 | 14.71 | 25.27 |
| **$a$19** | 4.21 | 2.37 | 5.36 |
| **$a$23** | 3.65 | 1.99 | 4.21 |
| **$a$27** | 3.22 | 2.59 | 3.59 |

**Table S2: Parameter estimates and their 95% confidence intervals for the wasp thermal performance curves.**

| Parameter | Estimate | 2.5% | 97.5% |
|---|---|---|---|
| **(Intercept)** | 1.83 | 1.44 | 2.22 |
| **G1** | -0.07 | -0.08 | -0.05 |
| **Factor: G0=23** | -1.51 | -2.04 | -0.97 |
| **Factor: G0=27** | -4.38 | -5.04 | -3.73 |
| **Factor: G1: G0=23** | 0.11 | 0.09 | 0.14 |
| **Factor: G1:G0=27** | 0.24 | 0.21 | 0.27 |

**Table S3: Description and values of parameters used in the simulation models.**

| Parameter | Description | Model A | Model B |
|---|---|---|---|
| $\boldsymbol{CT_{max}}$ | Fly critical thermal maximum (°C) | 30 | 30 |
| $\boldsymbol{K_H}$ | Fly carrying capacity | 100,000 | 100,000 |
| $\boldsymbol{f}$ | Proportion successful fly development | 1/20 | 1/20 |
| $\boldsymbol{h}$ | Wasp handling time | 0.01 | 0.01 |
| $\boldsymbol{t_p}$ | Maximum wasp attack coefficient | 0.000364 | 0.00026 |
| $\boldsymbol{k}$ | Wasp spatial aggregation term | NA | 0.65 |
| $\boldsymbol{d}$ | Wasp immigration | 10 | 10 |

**Table S4: Mean time to extinction, in generations, of the host-parasitoid system** depending on the inclusion of temporal autocorrelation in temperature and transgenerational effects of thermal conditions in the simulation model. Geometric mean with 100 runs per simulation scenario.

| *Temperature Autocorrelation* | *Transgenerational Effects Included* | | *Generations Until Extinction (Geometric Mean)* | |
|---|---|---|---|---|
| | **Fly** | **Wasp** | **Model A** | **Model B** |
| +0.8 | Yes | Yes | 967.72 | 1345.51 |
| +0.8 | Yes | No | 1372.47 | 1347.10 |
| +0.8 | No | No | 2103.27 | 2061.56 |
| +0.8 | No | Yes | 2103.04 | 2061.46 |
| 0 | Yes | Yes | 1351.81 | 1473.41 |
| 0 | Yes | No | 1492.65 | 1478.42 |
| 0 | No | No | 2298.41 | 2279.03 |
| 0 | No | Yes | 2298.39 | 2277.51 |

**Table S5 Supporting statistics for difference between simulation treatments under each model.** Statistical model formula: log10(Extinctions) ~ AC * FlyTrans * WaspTrans + (1 | TrialRep), where AC is a binary factor of autocorrelation, FlyTrans is a binary factor for the inclusion of fly transgenerational effects, and WaspTrans is a binary factor for the inclusion of wasp transgenerational effects. Per model there were 800 total observations (2x2x2 factors, 100 trials per treatment combination). Significance calculated via ANOVA using Satterthwaite's method.

| **Model** | **Term** | **Estimate** | **SE** | **Df** | **F-value** | **Pr(>F)** | **Significance** |
|---|---|---|---|---|---|---|---|
| **A** | (Intercept) | 3.3614282 | 0.0118 | | | | |
| | AC | -0.0385325 | 0.0165 | 693 | 61.62 | 0.0000 | * |
| | FlyTrans | -0.1874699 | 0.0165 | 693 | 814.15 | 0.0000 | * |
| | WaspTrans | -0.0000038 | 0.0165 | 693 | 34.95 | 0.0000 | * |
| | AC:FlyTrans | 0.0020767 | 0.0233 | 693 | 10.05 | 0.0016 | * |
| | AC:WaspTrans | -0.0000444 | 0.0233 | 693 | 10.89 | 0.0010 | * |
| | FlyTrans:WaspTrans | -0.0430403 | 0.0233 | 693 | 34.91 | 0.0000 | * |
| | AC:FlyTrans:WaspTrans | -0.1086623 | 0.0330 | 693 | 10.87 | 0.0010 | * |
| **B** | (Intercept) | 3.3577507 | 0.0029 | | | | |
| | AC | -0.0435547 | 0.0038 | 693 | 482.19 | 0.0000 | * |
| | FlyTrans | -0.1879517 | 0.0038 | 693 | 9689.53 | 0.0000 | * |
| | WaspTrans | -0.0002908 | 0.0038 | 693 | 0.0919 | 0.7619 | |
| | AC:FlyTrans | 0.0031547 | 0.0054 | 693 | 0.8510 | 0.3566 | |
| | AC:WaspTrans | 0.0002694 | 0.0054 | 693 | 0.0263 | 0.8711 | |
| | FlyTrans:WaspTrans | -0.0011846 | 0.0054 | 693 | 0.0488 | 0.8253 | |
| | AC:FlyTrans:WaspTrans | 0.0006931 | 0.0076 | 693 | 0.0083 | 0.9273 | |

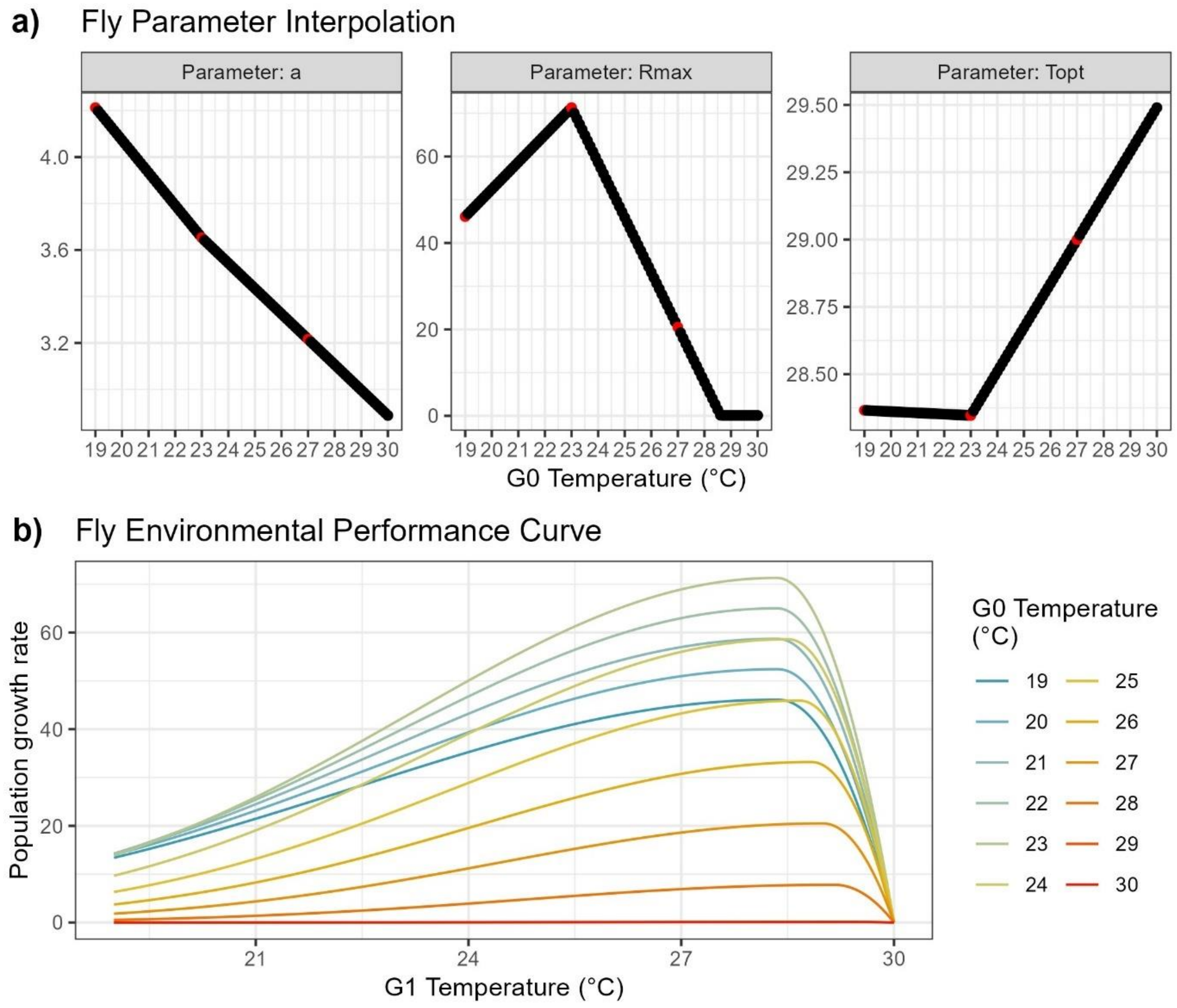


**Figure S1 Deriving fly population growth rate.** a) Values of the parameters $T_{opt}$, $R_{max}$ and $a$, depending on G0 temperature. Red points represent values at 19, 23 and 27°C, known from the best fit model. b) Population growth rate of the fly population, with the different colour lines representing different G0 temperatures.

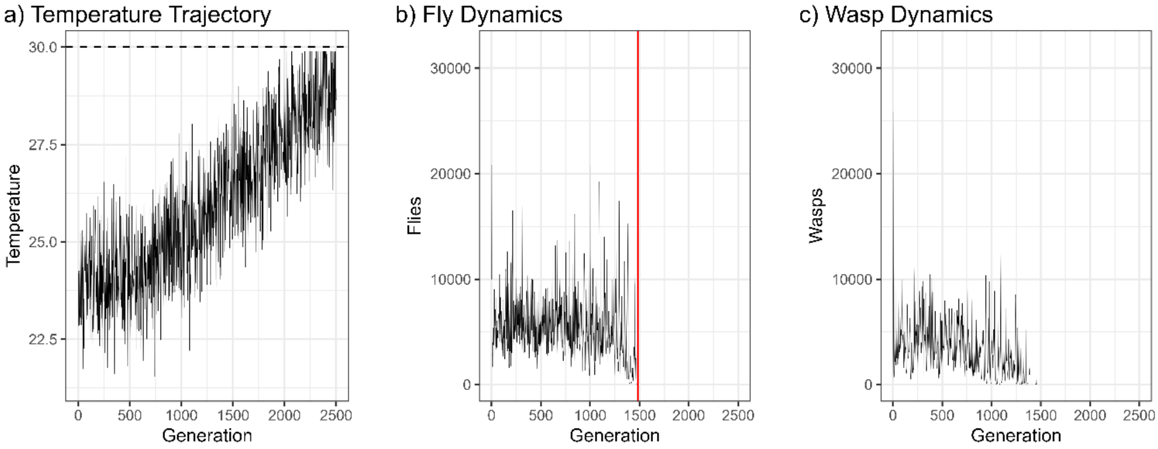


**Figure S2: Example time series from simulations.** a) Temperature trajectory with constant rate of underlying increase after the burn-in period, constant underlying variation and cap at 30 degrees, the critical temperature. b) Example trajectory of fly population dynamics, showing recorded extinction point in red. c) Corresponding example trajectory of wasp population dynamics.

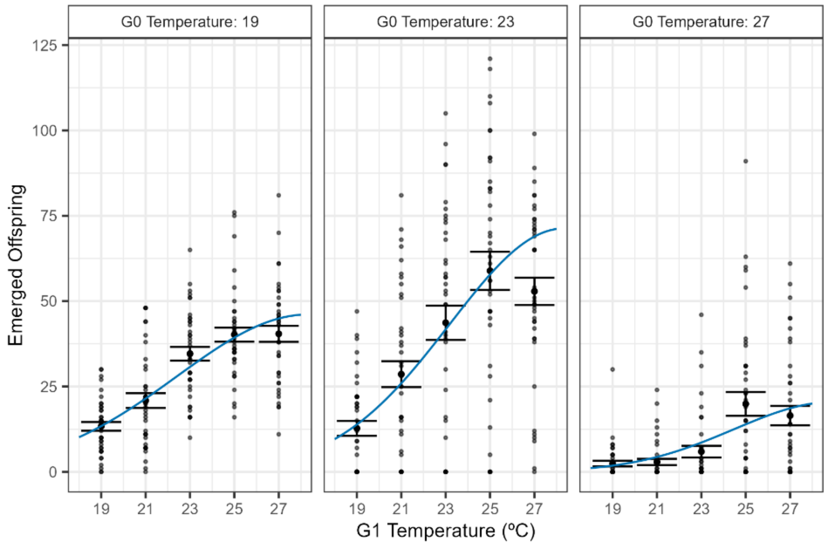


**Figure S3. Standard error around the mean for each temperature treatment combination.** Note that these errors were calculated assuming a Normal distribution, however the best fit thermal performance curves used a negative-binomial error model.

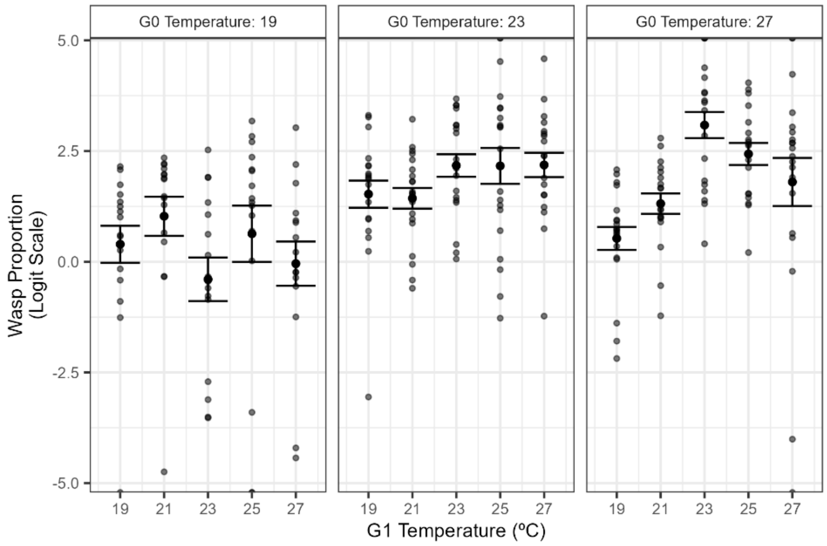


**Figure S4. Depiction of standard error around the mean proportion across the replicate vials for each temperature treatment combination.** Errors were calculated tased on the proportion within each vial on the logit scale. Each vial was weighted equally and vial proportions were upper and lower bounded at 0.01 and 0.99 to facilitate this presentation of the confidence of our data.
